# A Data-Driven Approach for the Development of a Time-informed Adverse Outcome Pathway-network for Cardiotoxicity of Environmental Chemicals

**DOI:** 10.64898/2025.12.22.695928

**Authors:** Alexandra Schaffert, Sivakumar Murugadoss, Tom Roos, Nunzia Linzalone, Gabriele Donzelli, Ronette Gehring, Birgit Mertens, Martin Paparella

**Affiliations:** Finnish Hub for Development and Validation of Integrated Approaches (FHAIVE), Faculty of Medicine and Health Technology, Tampere University, Tampere, Finland; Institute of Medical Biochemistry, Medical University Innsbruck, Innsbruck, Austria; Tampere Institute for Advanced Study, Tampere University, Tampere, Finland; Health Effects Laboratory, Department of Environmental Chemistry and Health Effects, NILU The Climate and Environmental Research Institute, Norway; Department of Population Health Sciences, Institute for Risk Assessment Sciences (IRAS), Faculty of Veterinary Medicine, Utrecht University, Utrecht, the Netherlands; Institute of Clinical Physiology of the National Research Council (CNR-IFC), Pisa, Italy; Scientific Department of Chemical and Physical Health Risks, Sciensano, Brussels, Belgium

**Keywords:** AOP network, key event relationship, essentiality, temporality, oxidative stress, mitochondrial dysfunction, cardiac remodelling, non-animal methods

## Abstract

We present a novel Adverse Outcome Pathway (AOP) network for environmental chemical-induced cardiotoxicity using a bottom-up, data-driven AOP development approach. Mechanistic endpoints were systematically extracted from 339 *in vitro* and *in vivo* studies, yielding 1,759 Key Event (KE) entries and 4,938 Key Event Relationship (KER) entries, including information on experimental methods, essentiality evidence (intervention experiments demonstrating upstream-downstream dependence), and study metadata. After quality filtering (high risk of bias, confounding cytotoxicity *in vitro*, excessive toxicity or animal well-being concerns *in vivo*, and low-frequency observations), 112 unique KEs and 829 unique KERs supported by at least three independent observations were retained for network construction. Network analysis identified oxidative stress and mitochondrial dysfunction as dominant hub processes linking diverse upstream perturbations to downstream cardiomyocyte injury, inflammation, cardiac remodelling (fibrosis and hypertrophy), decreased cardiac contractility, and reduced left ventricular function. Incorporating exposure duration at the KER level enabled time-resolved pathway interpretation and demonstrated that KE timing is relationship-dependent, revealing temporal patterns not apparent when analysing KEs in isolation. This evidence-weighted, time-resolved AOP network can support endpoint prioritisation and exposure-window selection for non-animal method (NAM) test batteries and mechanistically informed cardiotoxicity assessment.

**Synopsis:** Environmental chemicals converge on shared stress and injury pathways that drive cardiac remodelling and ventricular dysfunction. A time-resolved AOP network helps prioritise endpoints and exposure windows for non-animal cardiotoxicity testing.

## Introduction

Cardiovascular diseases (CVDs) remain the leading cause of mortality globally, accounting for a substantial fraction of all deaths each year [1]. Beyond established lifestyle and clinical risk factors, an expanding body of evidence indicates that environmental exposures, including industrial chemicals, complex mixtures, and ambient pollutants, contribute to cardiovascular morbidity [2, 3]. These exposures not only exacerbate pre-existing conditions like hypertension and heart failure but also contribute to the onset of cardiotoxicity [4]. Despite this growing concern, substantial knowledge gaps remain in identifying the specific mechanisms through which environmental chemicals impact cardiac function. These gaps present a significant barrier to the prediction and mitigation of chemical-induced cardiotoxicity, emphasizing the urgent need for a robust framework to address these challenges.

Existing regulatory frameworks for chemical safety assessment insufficiently assess cardiotoxicity [5]. For chemicals regulated under REACH, the Plant Protection Products Regulation, and the Biocidal Products Regulation, the evaluation of cardiotoxicity is not explicitly required. Instead, data on cardiotoxic effects are often derived indirectly from repeated-dose animal studies, which typically include endpoints such as cardiac weight, necropsy findings, and histopathology. While these studies provide some insight into potential cardiac toxicity, they do not capture detailed mechanistic information relevant to human health. Furthermore, the reliance on animal testing raises concerns about interspecies variability, resource demands, and alignment with the 3Rs (reduce, refine, replace) principles. These challenges highlight the need for more targeted and mechanistically informed approaches to assess cardiotoxicity in regulatory contexts.

Non-animal methods (NAMs), including human induced pluripotent stem cell-derived cardiomyocytes (hiPSC-CMs), organotypic cardiac models, high-content phenotyping, and multi-omics, offer a route to generate human-relevant mechanistic signals for cardiotoxicity, with the potential to increase throughput and improve biological interpretability. However, NAM outputs cannot be used optimally in isolation: regulatory application requires structured approaches that integrate diverse mechanistic and apical evidence into coherent causal narratives that support hazard identification, weight-of-evidence evaluation, and context-of-use decisions. Reflecting this need, cardiotoxicity is increasingly discussed as an emerging target for harmonisation and methodological development in international hazard assessment, including an OECD Working Party on Hazard Assessment (WPHA) initiative addressing cardiotoxicity as a potential hazard class and the role of novel methods in its evaluation [6, 7].

The Adverse Outcome Pathway (AOP) framework offers a structured approach to connect molecular-level perturbations caused by chemical exposures to higher-order adverse outcomes (AO), such as heart failure or reduced cardiac function [8]. AOPs rely on Key Events (KEs) and Key Event Relationships (KERs) to systematically link mechanistic data to observable AOs, thereby providing a transparent basis for mechanistically informed risk assessment. By integrating data generated with NAMs into AOPs, pathways leading to cardiotoxicity can be elucidated and critical points of intervention or regulatory relevance identified. Importantly, cardiotoxicity is unlikely to be captured by single linear pathways; rather, multiple converging and diverging processes can culminate in shared cardiac outcomes. AOP networks therefore provide an appropriate representation of mechanistic complexity and enable the identification of high-confidence, recurrent pathways across stressors and evidence streams [9]. Despite significant progress in AOP development for other toxicological endpoints, applications targeting cardiotoxicity remain scattered and often focus on individual pathways or pre-defined mechanisms, leaving a substantial gap in systematic, human-relevant AOP coverage for cardiac effects of environmental pollutants.

Here, we address this gap by implementing a strictly bottom-up, data-driven AOP-network development strategy for environmental chemical-induced cardiotoxicity. Building on our prior systematic evidence mapping (SEM) of pollutant-induced cardiotoxicity (Roos, Schaffert, and Murugadoss et al. under review at Environment International), we extracted reported mechanistic observations across *in vitro* and *in vivo* studies, harmonised these observations into candidate KEs and KERs, and assembled an evidence-weighted cardiotoxicity AOP network derived from the patterns emerging in the literature rather than from pre-defined pathways. We further introduce literature-derived temporality at the level of KERs, assigning time information to upstream-downstream event pairs, to enable time concordance assessment across the network and to better contextualise real-world applicability and AO development. The resulting network identifies the most prevalent and best-supported mechanistic trajectories from MIE to AO and provides a structured substrate for integrating NAM-derived evidence into cardiotoxicity hazard assessment and regulatory decision-making.

## Methods

### Overall AOP development strategy

The complete workflow used to translate heterogeneous cardiotoxicity literature into a consensus, time-resolved AOP is depicted in Figure 1. We began with a systematic review that identified 339 in-scope studies for mechanistic data extraction. For every study, two complementary data streams were captured: (i) study-level meta-information and (ii) AOP-relevant mechanistic information. Extraction was guided by a comprehensive KE library that combined cardiotoxicity-related KEs already listed in the AOP-Wiki with new, study-derived terms, and included AOs informed by previously established epidemiological findings. All entries then passed through a curation pipeline that harmonised KE terminology, applied quality and risk-of-bias filters, and normalised exposure times for AOP integration. The resulting high-confidence dataset was used to construct an evidence-weighted KER network, prioritised by essentiality support, empirical frequency, and KER-specific temporality. Expert review of biological plausibility and time concordance transformed this network into a consensus, time-resolved cardiotoxicity AOP.

**Figure 1:**
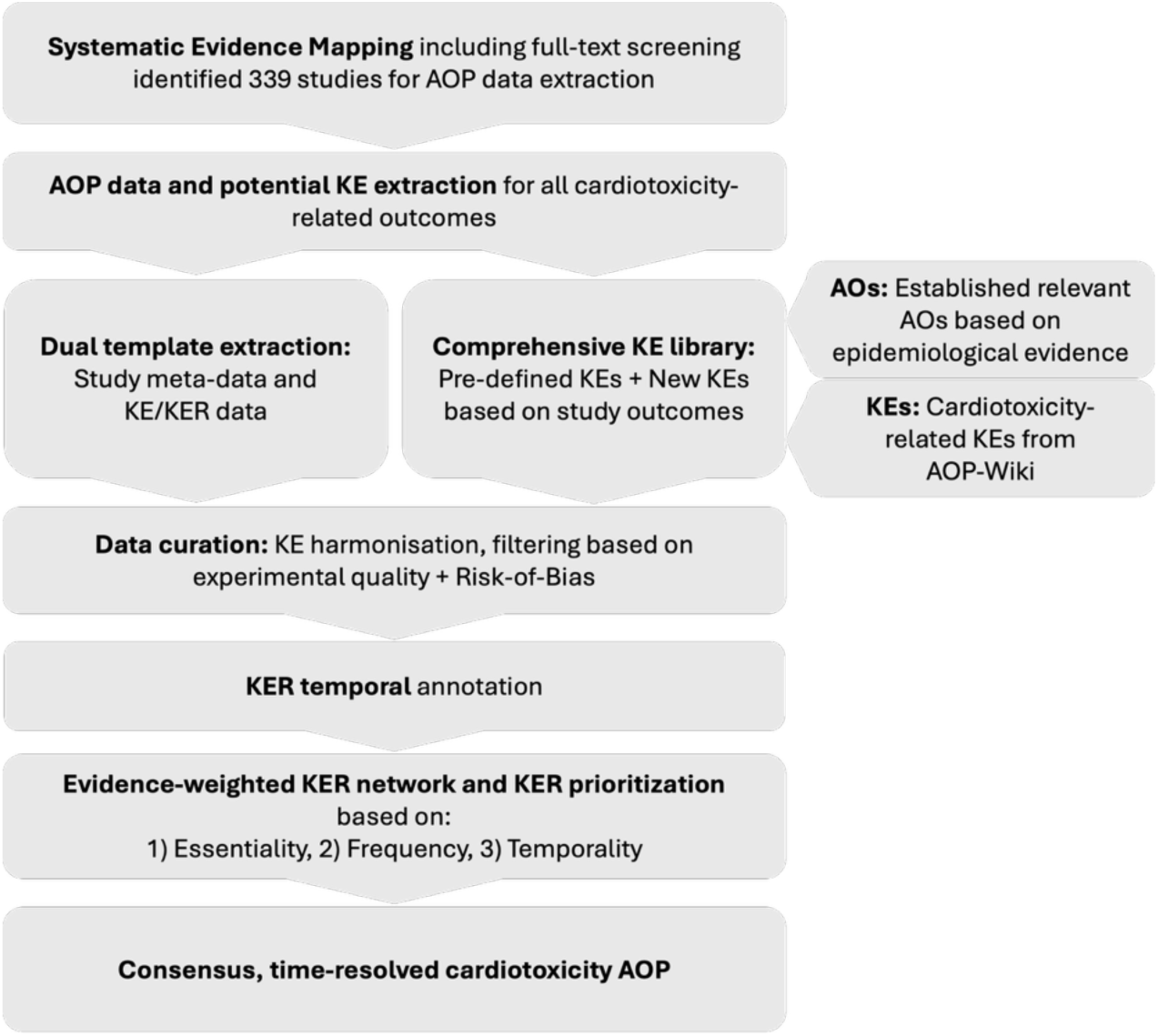
Data-driven AOP development workflow overview. From 339 studies, identified via a systematic review, potential KEs including new KEs and existing cardiotoxicity-related AOP-Wiki KEs, were extracted, followed by rigorous filtering, KE/exposure-time harmonisation, and evidence-weighted KER ranking. The process culminated in a consensus, time-resolved cardiotoxicity AOP network that integrates essentiality, empirical support, and KER-level temporality.

### Literature data extraction for AOP development

Literature data for the development of a cardiotoxicity AOP network was extracted from a systematic evidence mapping of studies related to environmental pollutant exposure leading to cardiotoxicity *in vitro* and *in vivo*, as previously described [10], (Roos, Schaffert, and Murugadoss et al. under review at Environment International). Extraction targeted all mechanistic and AO endpoints identified in the literature. Endpoints that had previously been identified in our epidemiological review of cardiotoxic effects associated with environmental chemical exposures in humans (Donzelli et al., 2024), were considered as potential AOs, and thus, terminology and definition criteria were harmonized to match those used in the epidemiological study. This ensured consistency in the annotation and clinical relevance of these outcomes within the AOP network.

Of the 400 studies included in the final systematic evidence mapping (under review at *Environment International*), 339 contained mechanistic information on at least two KEs and were therefore eligible for AOP-relevant data extraction, while the remaining 61 studies were excluded due to insufficient KE coverage. Relevant data were extracted into two initial templates. The first template included general study information, which encompassed details of study design, experimental setup, and a risk of bias (RoB) assessment conducted according to OHAT criteria (Table 1). The RoB assessment categorized studies into three tiers based on 11 questions, including two key questions, and was evaluated with a scoring system (Roos, Schaffert, and Murugadoss et al. under review at Environment International). The second template focused on AOP-relevant data, specifically targeting evidence to support the development of a cardiotoxicity AOP network (Table 1). Data from both extractions were made sure to align for consistency. For each study, key outcomes were extracted, alongside the methods used to assess these outcomes (e.g., qPCR), a description of outcomes (e.g., gene expression of pro-inflammatory cytokines was increased), and the corresponding KE (e.g., Inflammation). If multiple assays assessed the same KE within a study, only a single KE entry was recorded. In studies including both *in vitro* and *in vivo* experiments, KEs were extracted separately for each system. Only outcomes with statistically significant changes were considered as extractable KEs and no-effect outcomes excluded. This was noted as a limitation, as their absence precludes evaluation of potential opposing effects, although such reporting is generally limited in research and therefore represents a bias in itself. For each KE, it was recorded whether the outcome occurred only under (cyto)toxic doses or not (Table 1). In cases where essentiality of a KE was tested, such as, through the use of inhibitory agents targeting an upstream KE, the name of the upstream KE and the testing agent were recorded alongside the downstream KE. If a modulating factor (e.g., sex, diet, pre-existing conditions) significantly affected the outcome, this information was documented in relation to the corresponding KE.

**Table 1:** Overview of most relevant criteria extracted during the systematic review.

| Extraction item | Explanation |
| --- | --- |
| Biological system | Animal type, strain, life stage, or cell type |
| Chemical information | Chemical name, type, CAS number, purity, source, mixture components |
| Exposure scenario | Exposure duration, route, dose, number of concentrations tested |
| KE name | Best fitting name for potential cardiotoxicity-related KE (harmonized among data) |
| Essentiality assessment | Essentiality of a KE and how it was tested, e.g., addition of antioxidant which inhibits KE "Increase oxidative stress" lead to diminishing of downstream KE "Increase cell death/cell injury". Thus, during extraction of KE "Increase cell death/cell injury", essentiality testing was entered in the row and "Increase oxidative stress" was entered as upstream connected essential KE. |
| (Cyto)toxicity | *In vitro*: Yes/No for significant cytotoxicity at KE-relevant dose, or if cytotoxicity was not tested<br>*In vivo*: Yes/No for reported cases of mortality or significant decrease in body weight compared to the control, or if body weight was not reported at the end of experiment |
| Modulating factors | Factors which affected the intensity of the KE (e.g., sex, diet, preexisting conditions) |
| Method/assay | Experimental method or assay which was used to assess the KE |
| Outcome | Description of the outcome associated with the KE |
| Risk of Bias assessment | Risk of Bias was assessed for each study, consisting of 11 questions, including 2 key questions. |

In the first extraction round, mechanistic outcomes were mapped to an initial controlled vocabulary of predefined KE terms derived from cardiotoxicity-relevant AOP-Wiki entries and to AO definitions aligned with epidemiological evidence. Where none of the predefined KEs adequately captured the reported effect, extractors assigned new candidate KE terms directly from the study-derived observation. Importantly, this procedure resulted in an empirically expanded KE library in which the majority of KEs captured in the evidence base were not covered by the predefined AOP-Wiki-derived terms. The predefined KE vocabulary was intentionally restricted to three cardiotoxicity-relevant AOP-Wiki entries: AOP 433 [11], AOP 261 [12], and 262 [13]; broader cardiovascular AOPs were not included because they fell outside the defined study scope. Following consolidation and harmonisation of terminology across all extracted observations, a second extraction round re-mapped all KE annotations using the standardised KE set to ensure consistency and reduce synonym-driven fragmentation. AOP data extraction was performed equally by three authors (A.S., S.M., and T.R.) and cross-checked by A.S. and S.M. Consistency and extraction quality were evaluated independently by the authors, yielding a curated, quality-controlled dataset comprising AOP-specific mechanistic evidence linked to study-level metadata.

### Data curation, processing, and evaluation

Following data extraction, all entries underwent manual curation to ensure consistency in KE assignments and to correct any extraction-related errors. Subsequent data processing was performed in R Studio (version 4.4.1). AOP-specific information was integrated with general study-level metadata, and a quality filtering step was applied to exclude entries that did not meet predefined criteria. This included removal of Tier 3 studies based on the risk of bias assessment, *in vitro* studies associated with cytotoxic concentrations (as assessed within the respective studies), and *in vivo* studies reporting excessive body weight loss or mortality.

To improve interpretability and reduce redundancy, closely related KEs were consolidated based on mechanistic pathway evidence. This included the grouping of multiple signaling events associated with cardiac hypertrophy or fibrosis under unified KE descriptors reflecting their shared biological function.

For the construction of KERs, KE pairs were derived from the co-occurrence of two KEs within the same study experiment, representing “candidate relationships” that require biological plausibility and (where available) essentiality to support directionality. Directionality was assigned where experimental data supported biological essentiality, that is, where perturbation of an upstream KE (e.g., through inhibition or genetic manipulation) resulted in a measurable change in the downstream KE. Only quality-filtered data were used for all subsequent analyses, including network construction and interpretation.

### KER time annotation

To characterise the temporal progression of cardiotoxic effects, we first assigned an exposure duration to each KE-study record based on the shortest duration at which a statistically significant modulation of that KE was reported. If a study reported only one time point, that duration was used. If multiple time points were assessed, we extracted the earliest time point at which the KE showed a statistically significant change compared with control (as defined by the study authors). Free-text descriptions of exposure schedules were parsed and converted into numeric exposure durations in days. Because the dataset covered multiple species and test systems with different typical study durations, we harmonised exposure times using species- and system-specific cut-offs anchored in OECD Test Guidelines and common toxicology study designs. These cut-offs were defined a priori for each experiment type (*in vivo* vs. *in vitro*) and for each species category (mice, rats, zebrafish, Drosophila, swine, chickens, rabbits, quails, tadpoles, marmosets, and *in vitro* systems). Exact thresholds and supporting references are summarised in Supplement 1, Table S1. Each KE-study record was then assigned to one of four initial duration categories: Acute, Subacute, Subchronic, or Chronic. Briefly, for *in vivo* mammalian and avian studies, Acute exposure was defined as a single administration with a total duration below the species-specific acute threshold (typically up to 1 day). Dosing regimen was inferred from the study description. Repeated or continuous dosing, even if shorter than the acute threshold, was therefore classified as Subacute. Subacute, Subchronic, and Chronic categories covered increasingly longer exposure windows up to and beyond the species-specific cut-offs corresponding to subacute (for example 28 days in rodents), subchronic (for example 90 days in rodents), and chronic exposures (multi-month or lifetime), respectively. For aquatic and invertebrate *in vivo* models (zebrafish, tadpoles, Drosophila), acute windows were adjusted to reflect shorter life cycles and typical guideline practice. Continuous exposures of a few days were still classified as Acute when they remained below the species-specific acute threshold; longer exposures were assigned to Subacute, Subchronic, or Chronic according to the cut-offs listed in Supplementary Table S1. For *in vitro* studies, exposure duration classification was based solely on incubation time, independent of dosing frequency. Exposures were categorised as Acute (up to 3 days), Subacute (more than 3–14 days), Subchronic (more than 14–28 days), or Chronic (more than 28 days), reflecting common practice in mechanistic and omics-based assays. For descriptive summaries and network visualisation, Subchronic and Chronic categories were combined into a single Chronic class, because explicitly chronic studies were overall rare (Supplement 1, Figure S2) and both subchronic and chronic designs represent sustained perturbations compared with shorter-term exposures. The full category classification and all species-specific thresholds were documented in Supplementary Table S1.

All duration categories were assigned from the reported exposure times using species-specific time windows derived from typical test guideline durations and used as heuristic anchors. Exposure durations were not adjusted for species differences in internal dose or accumulation. Thus, the categories represent a structured grouping of study designs rather than harmonised internal exposure conditions.

### AOP network development

After systematic extraction, quality-filtering and curation, KERs were further refined based on biological plausibility. Specifically, KERs directly linking molecular-level KEs to organ- or organism-level KEs were excluded, as they bypass intermediate cellular and tissue-level events that are required for mechanistically interpretable pathways. All remaining KERs were aggregated into an evidence-weighted KE–KE network, retaining only those KERs that were supported by at least three independent study observations. This network was then evaluated in a three-step evidence-weighting procedure: 1) Empirical support: the number of independent KER observations; 2) Essentiality support: KERs underpinned by targeted perturbation experiments were treated as directed, high-confidence edges; 3) KER-specific temporality: for each KER, the shortest reported exposure duration associated with the occurrence of the downstream KE was assigned, enabling a network-wide assessment of time concordance. Due to substantial heterogeneity in chemical identity, experimental design and dose selection across the included studies, dose–response information could not be meaningfully harmonised; dose concordance analysis was therefore not performed. The final product is a consensus, time-resolved cardiotoxicity AOP network that captures the most empirically supported and temporally coherent pathways emerging from the evidence map.

## Results

### Experimental model systems captured in AOP evidence mapping

Across the extracted dataset, *in vivo* evidence contributed the majority of KE entries, with *in vitro* studies representing a smaller but methodologically diverse fraction (Figure 2A). Within the *in vivo* evidence stream, rodent models dominated, with rats most frequently represented followed by mice; all other species (e.g., chicken, quails, zebrafish, swine/pigs, rabbits, Drosophila, tadpoles, and marmosets) contributed comparatively fewer KE entries (Figure 2B). *In vitro* KE entries were distributed across a broad set of cardiac-relevant cellular systems, led by primary cardiomyocytes, H9c2 cells, and human cardiomyocytes, with additional coverage from endothelial and fibroblast models (e.g., EA.hy926, HUVEC, neonatal cardiac fibroblasts) and stem cell-derived platforms (e.g., hiPSC- and mESC-derived cardiomyocytes) (Figure 2C). Collectively, this model landscape indicates that the mechanistic evidence base used for bottom-up AOP-network construction is anchored in mammalian *in vivo* biology, while simultaneously capturing human-relevant cellular models that are directly compatible with NAM-oriented mechanistic testing and provide coverage for multi-cell-type processes (e.g., vascular/endothelial contribution and fibrosis-related mechanisms).

**Figure 2:**
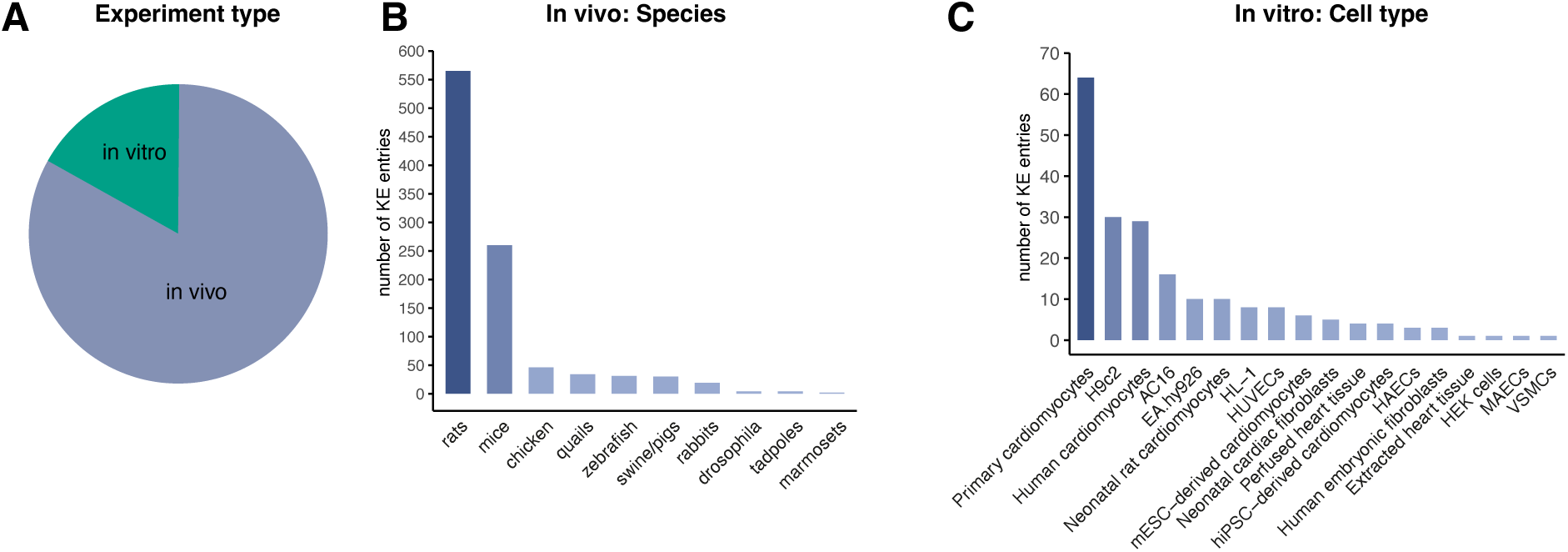
Biological models used in studies included for AOP extraction. **(A)** Proportion of KE entries derived from in vivo versus in vitro studies. **(B)** Species distribution in in vivo KE entries. **(C)** Cell types used in in vitro KE entries.

### Network essentiality analysis reveals central role of oxidative stress in cardiotoxicity of environmental chemicals

From 339 studies included in the extraction, 1,759 KE entries and 4,938 KER entries were retrieved (Figure 3). A quality filtering step excluded entries due to high risk of bias, significant cytotoxicity (*in vitro*), signs of excessive toxicity (*in vivo*), or low frequency, to ensure data robustness and improve confidence in the KE/KERs. Applying the evidence threshold for network inclusion (≥3 independent observations per relationship) yielded 112 unique KEs and 829 unique KERs, which formed the basis for subsequent network construction.

**Figure 3:**
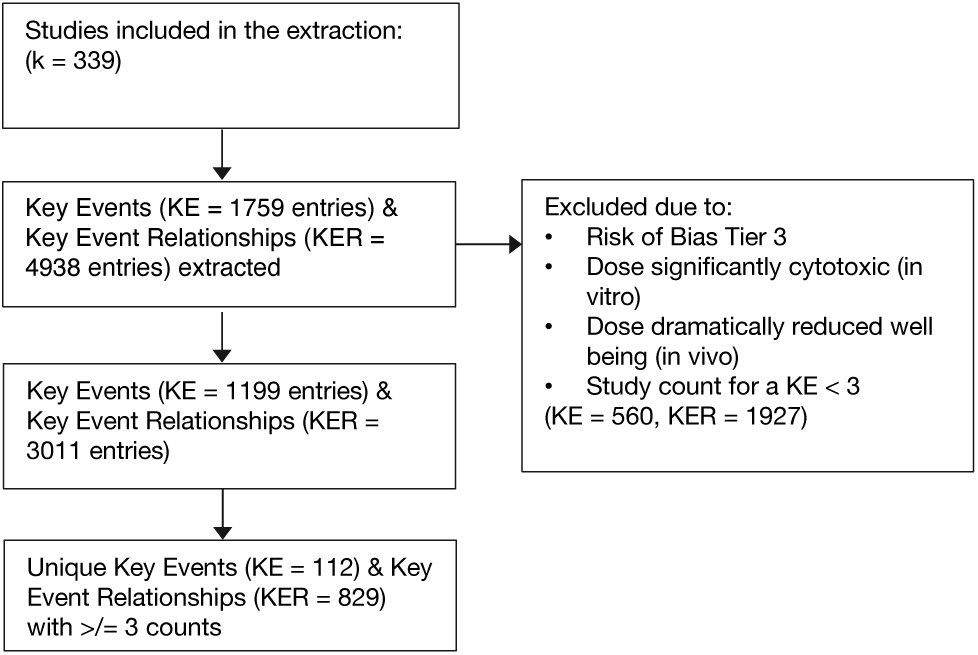
Flow chart of AOP data extraction during literature review.

To prioritise mechanistically substantiated relationships and support directionality, we constructed an essentiality-informed network based on intervention evidence. Essentiality-supported KERs were defined as relationships in which targeted perturbation of an upstream KE (e.g., pharmacological inhibition or genetic manipulation) resulted in a measurable change in a downstream KE, and only relationships observed at least three times were retained (Figure 4).

**Figure 4:**
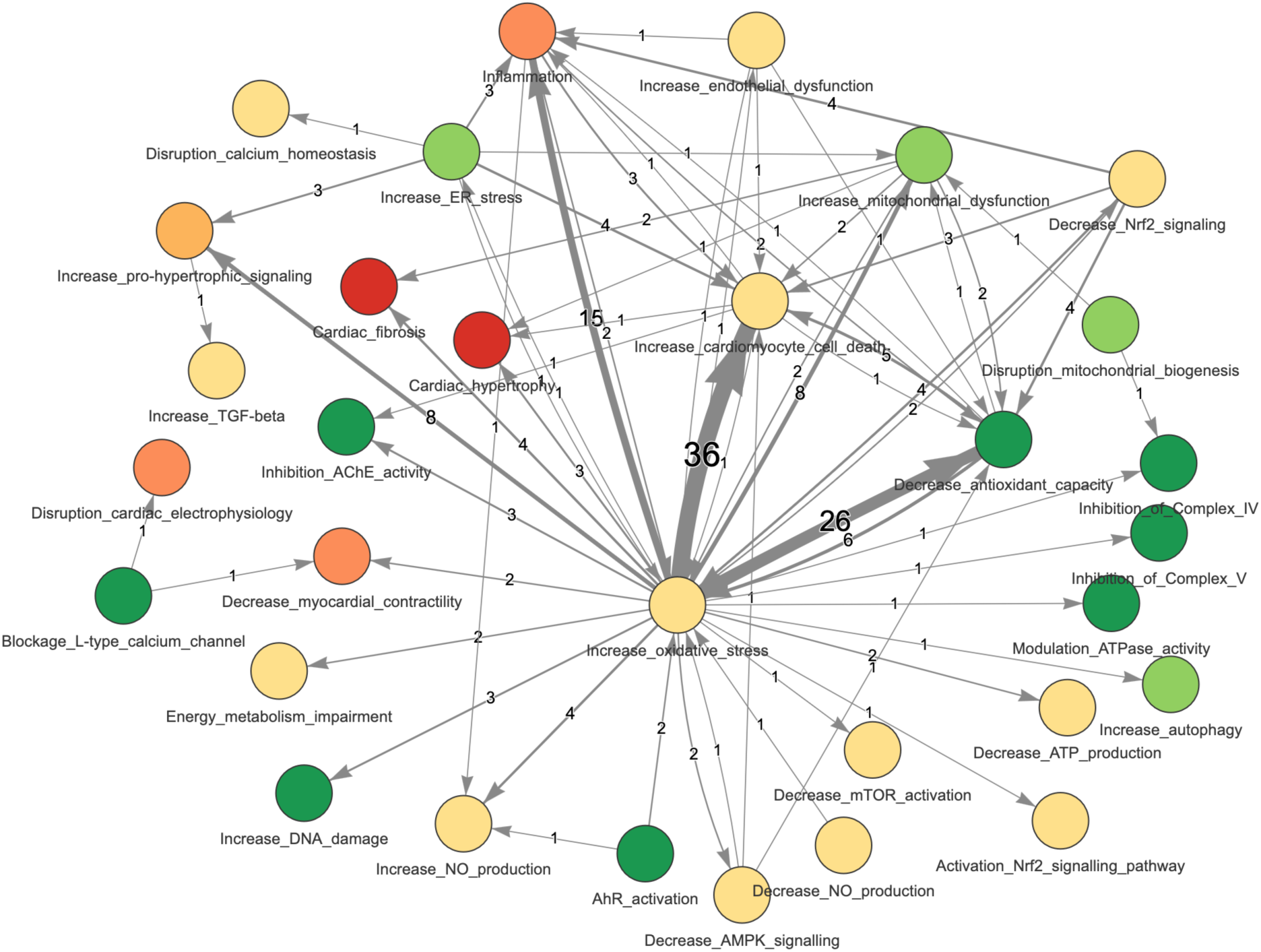
Network of extracted essentiality relationships. Edges represented essentiality relationships, with their thickness proportional to the essentiality count. Nodes represent KEs and are color-coded based on their biological level (dark green: Molecular, light green: Sub-cellular; yellow: cellular; light orange: multi-cellular; orange: tissue; red: organ/organism).

The essentiality network (Figure 4) highlights the central role of oxidative stress in cardiotoxicity, serving as a major hub that connects multiple events across different biological levels. *Increase oxidative stress* exhibited the highest essentiality count, with strong, experimentally supported links to *increase cardiomyocyte cell death* (36 relationships), *decrease antioxidant capacity* (n=26), and *inflammation* (n=15). Other well-supported relationships include connections to *mitochondrial dysfunction* and *pro-hypertrophic signaling* (n=8 each), and to *cardiac fibrosis* (n=4). While oxidative stress was the most interconnected node, bidirectional interactions were also observed. For example, *decreased antioxidant capacity* emerged both as a downstream consequence and as an upstream driver of oxidative stress. Similarly, *mitochondrial dysfunction* and *inflammation* were reciprocally linked to oxidative stress, reinforcing its centrality in the network.

The presence of Aryl Hydrocarbon Receptor (AhR) activation within the network, although with fewer essentiality-supported relationships, suggests that this MIE may act through modulation of oxidative stress and its downstream consequences. Overall, the essentiality-informed structure underscores oxidative stress as a mechanistically validated and central driver of cardiotoxic responses, while also illustrating how multiple feedback loops and intermediate signals shape the broader biological landscape. The presence of *AhR activation* within the network, albeit with fewer essentiality-supported relationships, suggests that this MIE may act through modulation of oxidative stress and its downstream consequences. Overall, the essentiality-informed structure underscores oxidative stress as a mechanistically validated and central driver of cardiotoxic responses, while also illustrating how multiple feedback loops and intermediate signals shape the broader biological landscape.

### Integrating network evidence and temporal progression

To contextualise network structure in time, we assessed whether exposure-duration categories exhibited a consistent monotonic pattern across biological levels (MIE → intermediate KEs → AOs). No generalisable trend was observed at the level of biological level categories (Supplement 1, Figure S2), indicating that “level” alone does not reliably encode temporal progression in the extracted evidence base. We found that for each KE, observations are distributed across acute, sub-acute, and chronic categories depending on which specific KER it participates in (i.e., which KE it co-occurred with), as exemplified by the hub KE increase oxidative stress (Figure 5A). In other words, the same KE does not have a single characteristic “time“, but its temporal context shifts with the relationship in which it is embedded. We therefore evaluated temporality at the KER level. Accordingly, we assigned each KE pair the most frequently observed exposure-duration category of both connected KEs, yielding a KER-specific temporal annotation that can be propagated across the network to visualise time-resolved mechanistic trajectories (Figure 5B–D). In each subnetwork, edge thickness reflects empirical support (number of independent observations for the respective KER within the evidence base), while arrow direction indicates essentiality-supported directionality (i.e., intervention evidence demonstrating that perturbation of an upstream KE modifies a downstream KE). This representation allows visual identification of (i) high-frequency KERs within each duration window and (ii) where directionality is mechanistically substantiated, supporting causal ordering within frequently reported pathways.

**Figure 5:**
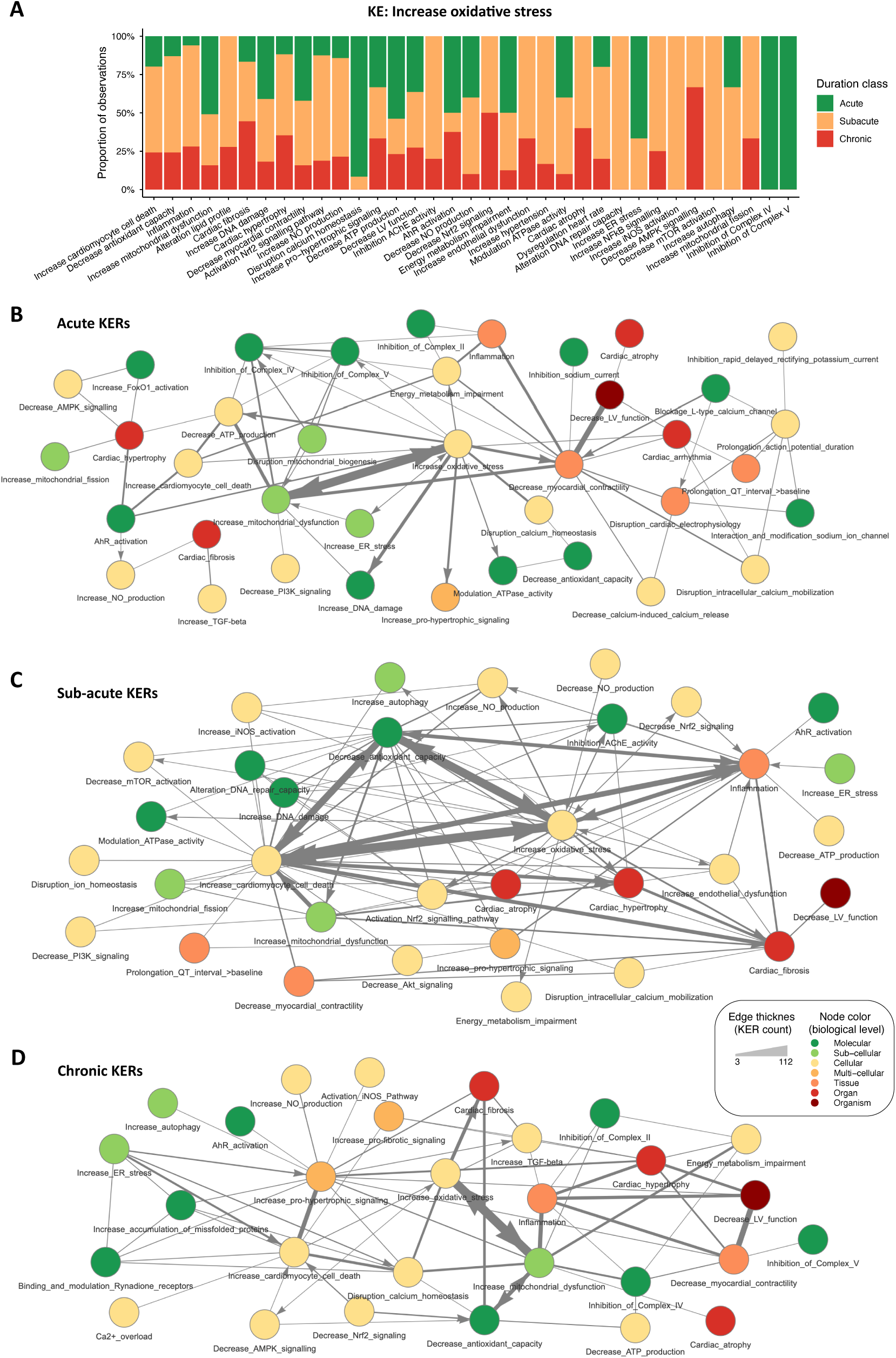
Duration-organised, evidence-weighted cardiotoxicity adverse outcome pathway (AOP) subnetworks and Key Event Relationship (KER)-specific temporality for a representative hub Key Event (KE). (A) Example KE “Increase oxidative stress” showing for each co-occurrence with a connected KE (x-axis), the proportion of observations assigned to acute, sub-acute, or chronic exposure-duration classes (y-axis), illustrating that the same KE can occur across different exposure windows depending on the specific upstream/downstream relationship. (B–D) Subnetworks of KERs assigned to (B) acute, (C) sub-acute, and (D) chronic duration classes. Nodes represent KEs and are colour-coded by biological level (legend). Edge thickness is proportional to empirical support (KER count).

Notably, *in vitro*-derived KE entries were predominantly acute (Supplement 1, Figure S1), reflecting typical short *in vitro* exposure windows; longer *in vitro* designs occurred but were uncommon, and this bias is unlikely to affect overall patterns because *in vitro* entries represent a minority of the dataset (Figure 2A).

### Acute subnetwork: Early functional perturbation converging on contractility and electrophysiology

In the acute subnetwork (Figure 5B), the network is characterised by a sparse set of predominantly low-supported electrophysiology- and calcium-handling-related KERs. These include ion-channel and action potential-related events that link to disruption of cardiac electrophysiology, prolongation of action potential or QT interval, and *cardiac arrhythmia*. Although these relationships are present, they are supported by relatively few observations and form peripheral structures rather than dominant network hubs. Nevertheless, these electrophysiological perturbations connect to *decrease myocardial contractility*, placing acute electrical disruption in proximity to early functional impairment.

In parallel, molecular and subcellular perturbations are already evident in the acute window. Inhibition of mitochondrial respiratory chain complexes and *AhR activation* are connected to intermediate cellular stress responses, including *increase oxidative stress*, *increase mitochondrial dysfunction*, *decrease ATP production*, and energy metabolism impairment. These stress- and energy-related KEs show stronger connectivity than electrophysiological events and converge on *decrease myocardial contractility*, which emerges as a central functional outcome in the acute network. Together, the acute subnetwork indicates that early cardiotoxic effects are driven primarily by rapid metabolic and stress-related perturbations, with electrophysiological alterations appearing as additional but less consistently supported contributors.

### Sub-acute subnetwork: Shift toward injury, antioxidant capacity, and inflammation

In the sub-acute subnetwork (Figure 5C), network structure shifts toward cellular injury and inflammatory processes as dominant organising features. *Increase cardiomyocyte cell death* emerges as a central hub with multiple high-support incoming and outgoing relationships. Strongly connected upstream KEs include *increase oxidative stress*, *decrease antioxidant capacity*, and *increase mitochondrial dysfunction*, indicating consolidation of stress-related pathways into cellular injury. Downstream, *increase cardiomyocyte cell death* links to tissue- and organ-level outcomes, including *cardiac fibrosis* and *cardiac hypertrophy*.

*Inflammation* becomes a second highly connected node in the sub-acute network. It receives inputs from oxidative stress and cell injury and connects onward to both fibrotic and hypertrophic pathways. Compared with the acute subnetwork, the sub-acute network contains a richer set of intermediate regulatory KEs, including mitochondrial dynamics (e.g., mitochondrial fission), stress-response signalling (e.g., Nrf2-related changes), and homeostatic processes such as autophagy and mTOR signalling. Overall, the sub-acute subnetwork marks a transition from early functional disturbance to accumulating cellular damage and inflammatory signalling that mechanistically links initial stress responses to emerging structural remodelling.

### Chronic subnetwork: Sustained stress-mitochondrial dysfunction coupled to remodelling and LV function decrease

In the chronic subnetwork (Figure 5D), network structure becomes more compact and is dominated by strongly supported KERs centred on *increase oxidative stress*, *decrease antioxidant capacity*, and *increase mitochondrial dysfunction*. These KEs form a persistent stress-mitochondrial dysfunction axis that anchors the chronic network and connects extensively to downstream inflammatory and remodelling-related processes. *Cardiac fibrosis* emerges as a key tissue-level outcome in the chronic network, linked to upstream oxidative stress and connected to pro-fibrotic signalling, including *increase TGF-beta*. *Cardiac hypertrophy* is similarly integrated, connecting to *increase pro-hypertrophic signalling* and *inflammation*, and linking onward to functional impairment. *Decrease myocardial contractility* and *decrease LV function* are now tightly embedded within the chronic network, receiving inputs from sustained stress, mitochondrial dysfunction, inflammation, and structural remodelling pathways. Calcium-related disturbances and *increase ER stress* primarily connect through injury-related endpoints such as cardiomyocyte cell death rather than acting as dominant organising features. Collectively, the chronic subnetwork indicates that long-term cardiotoxicity is driven by persistent oxidative and mitochondrial stress, leading to inflammatory amplification, structural remodelling, and progressive decline in ventricular function.

### Modulating factors and potential prototypical stressors

Several modulating factors were reported to influence the occurrence or magnitude of cardiotoxic KEs (Figure 6A). High-fat diet was the most frequently reported modifier and was associated with amplification of structural, metabolic, and inflammatory KEs, including *cardiac hypertrophy*, *energy metabolism impairment*, and *inflammation*. Pre-existing cardiovascular conditions similarly intensified downstream KEs, particularly *increase oxidative stress*, cell injury/death, and *cardiac fibrosis*. Sex-specific effects were observed in a small number of studies, mainly affecting cell injury/death and cardiac remodelling KEs, but the limited evidence precludes generalisation. Overall, modulating-factor information was inconsistently reported, highlighting substantial gaps in contextual data relevant to cardiotoxicity susceptibility.

**Figure 6:**
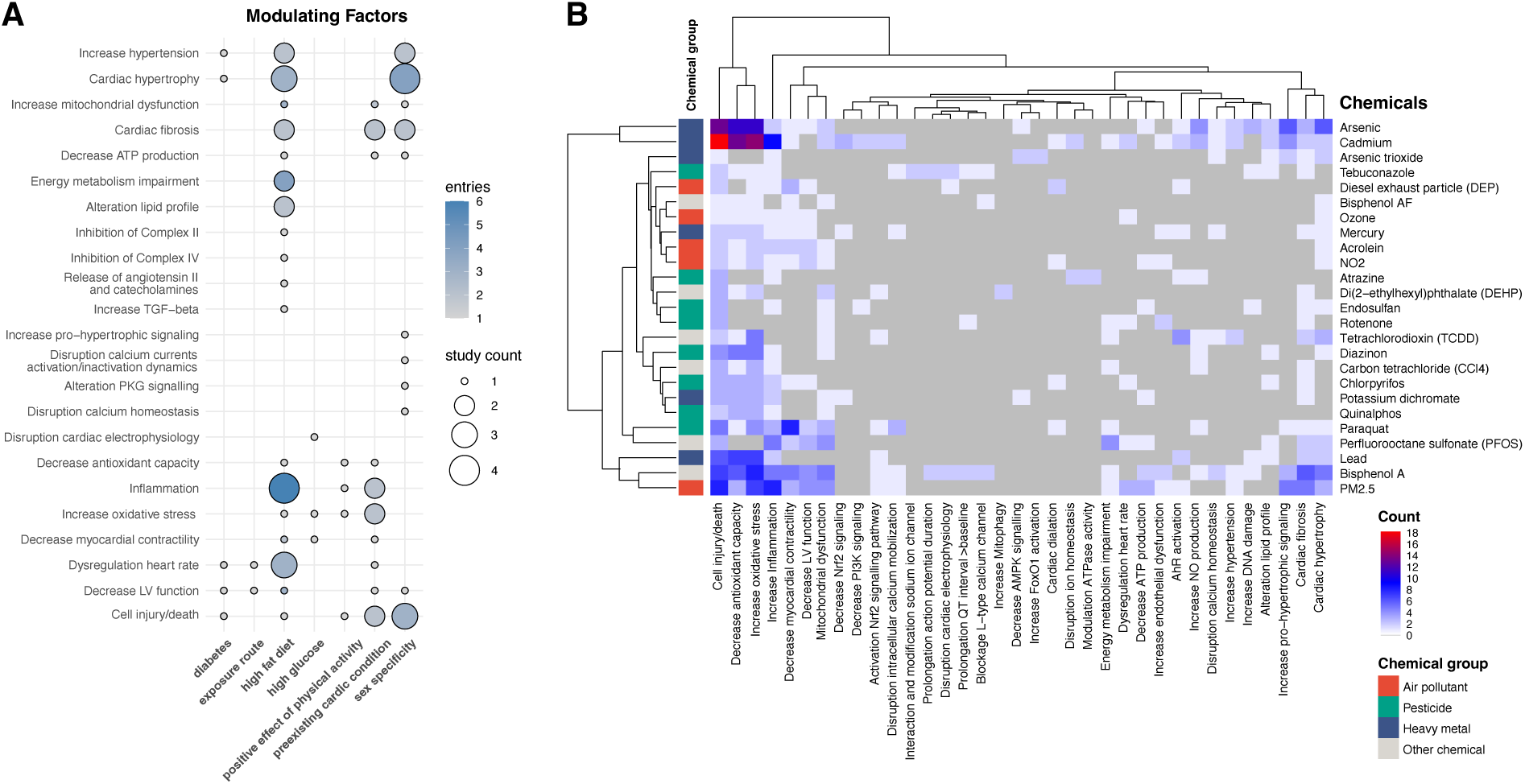
Modulating factors and chemical-KE associations. (A) Bubble plot showing seven modulating-factor categories that influenced cardiotoxic KEs. (B) Heat-map of chemicals with top associations, based on chemicals found with at least 10 associations, and KEs with at least 4 associations. Chemical group and number of studies that investigated the chemicals are indicated on the right side.

To explore whether specific environmental chemicals preferentially associate with particular KEs in the cardiotoxicity AOP network, we analysed stressor–KE co-occurrence patterns for chemicals with sufficient representation in the dataset (Figure 6B). Across chemicals, *increase oxidative stress, inflammation, and increase mitochondrial dysfunction* emerged as the most frequently reported KEs, consistent with their central positioning in the network analysis. Several stressors showed recurrent associations with these pathways rather than with highly specific downstream outcomes. In particular, the heavy metals cadmium and arsenic were strongly associated with the MIE *decrease antioxidant capacity* and with downstream KEs including *increase oxidative stress* and *cell injury/death*. Lead exhibited a similar association pattern, albeit supported by fewer studies, resulting in lower overall evidence density. Cadmium additionally showed a comparatively strong association with *inflammation*, reinforcing its link to stress- and injury-driven cardiotoxic mechanisms. The persistent organic pollutant 2,3,7,8-tetrachlorodibenzo-p-dioxin (TCDD) was frequently associated with *AhR activation* and *increase oxidative stress*, reflecting its well-characterised mode of action and supporting the inclusion of *AhR activation* as a molecular-level entry point into the cardiotoxicity network. Across the dataset, however, little clustering of chemicals by KE profiles was observed. This lack of clear chemical grouping likely reflects a combination of limited data availability for individual chemicals, substantial heterogeneity in study designs and endpoints, and strong imbalances in the number of studies per stressor. As a result, the extracted evidence preferentially supports mechanistic convergence on shared stress and injury pathways, rather than stressor-specific, KE-distinct cardiotoxic signatures.

## Discussion

Despite increasing recognition of environmental contributors to CVD, cardiotoxicity remains poorly represented in current AOP resources and is rarely addressed through systematic, pathway-resolved hazard frameworks. Here, we address this gap by applying a strictly bottom- up, literature-driven AOP development strategy to derive a mechanistically structured and time-resolved cardiotoxicity AOP network for environmental chemicals. By coupling systematic evidence mapping with evidence-weighted network assembly and KER-level temporal annotation, we provide a high-confidence synthesis of the cardiotoxicity mechanisms most consistently supported across heterogeneous *in vitro* and *in vivo* studies, together with an interpretable temporal context for sequence-of-events reasoning.

Our framework is grounded in evidence extracted from 339 studies identified via systematic evidence mapping of pollutant-induced cardiotoxicity. After quality filtering, the final network construction was based on 112 harmonised KEs and 829 unique KERs supported by at least three independent observations. Directionality was assigned where essentiality evidence was available (i.e., targeted perturbation of an upstream KE altered a downstream KE), and exposure-time information was harmonised across species and test systems to enable categorisation of KERs into acute, sub-acute, and chronic windows. The resulting network integrates three complementary evidence layers: empirical support, mechanistic directionality (where available), and temporal context. This combination produces an AOP network that is evidence-weighted and directly interpretable for NAM-oriented testing strategies, where the selection of endpoints and exposure durations is a key design variable.

### Central mechanistic spine and convergence across stressors

Data-driven assessment of KER frequency and essentiality highlighted oxidative stress and mitochondrial dysfunction as the most consistently represented events. This prominence likely reflects both their central role in cardiotoxic mechanisms and the relative sensitivity of current experimental models to detect these pathways. These events are not only biologically plausible but are also commonly investigated endpoints in both *in vitro* and *in vivo* systems, which may partially explain their strong representation in the dataset. Their frequent co-occurrence and essentiality-supported relationships across studies support their inclusion as central components in an environmental chemical-induced cardiotoxicity AOP network. Mitochondria are vital for cardiac function, supplying the bulk of myocardial ATP while acting as central redox hubs; their impairment simultaneously depletes cellular energy and elevates ROS. Whether oxidative stress precedes mitochondrial dysfunction or arises secondarily remains debated, but our KER-level temporal labels show that both events can occur within the same acute exposure window, consistent with a bidirectional feed-forward loop shown in the essentiality evidence network. Thereafter, pro-hypertrophic signalling and inflammatory amplification appear predominantly in the sub-chronic and chronic categories, leading to structural remodelling (cardiac fibrosis, hypertrophy) and functional decline (decreased LV function). Mitochondrial dysfunction and oxidative stress as key drivers of AOs such as cardiac remodelling and LV dysfunction align well with established literature knowledge [14–16].

The final network structure also clarifies how this stress-centric spine progresses toward AOs. Molecular/subcellular perturbations (e.g., inhibition of mitochondrial electron transport chain complexes, direct redox-mediated ROS production, and AhR activation) converge on oxidative stress and mitochondrial dysfunction within acute exposure windows. From there, the dominant progression is routed through cell injury/death and inflammation in sub-acute windows, followed by chronic engagement of pro-hypertrophic signalling and remodelling processes. This culminates in cardiac hypertrophy and cardiac fibrosis (captured as remodelling outcomes), and ultimately decreased LV function. This trajectory provides a coherent, evidence-weighted mechanistic narrative for environmental cardiotoxicity that aligns with the predominant endpoint types observed in the evidence base (stress and injury pathways leading to remodelling and functional decline).

While the network captures many intermediate drivers of cardiotoxicity with high evidence, few MIEs were robustly supported. The most frequently observed MIE was inhibition of mitochondrial electron transport chain complexes (I–V; strongest for complex IV), followed by AhR activation and direct redox-related ROS generation. Other proposed MIEs (e.g., receptor-mediated signalling, metabolic disruption, electrophilic covalent binding) were rarely reported or absent, likely reflecting a literature focus on downstream phenotypes. Given growing recommendations to recognise cardiotoxicity as a standalone regulatory endpoint [7], future studies should more systematically identify and report diverse MIEs to strengthen AOP coverage and support cardio-specific NAM batteries.

### Alignment with epidemiology and adverse outcomes of regulatory relevance

The prominence of structural cardiac effects/cardiac remodeling (including hypertrophy and fibrosis) and LV dysfunction as AOs in our data-driven AOP network closely aligns with evidence from recent epidemiological research. A systematic review and meta-analysis of human observational studies demonstrated that exposure to air pollutants (e.g., PM₂.₅, PM₁₀, NO₂) and heavy metals is associated with early abnormalities in myocardial structure and function, including both systolic and diastolic LV dysfunction [17]. Notably, the meta-analysis of epidemiological data highlights these endpoints as early markers of heart failure risk and recommends their inclusion in future longitudinal and case-control studies. This concordance strengthens the biological plausibility and translational value of our network, and illustrates how mechanistic AOP frameworks can complement population-based evidence to confirm causal relationships between environmental chemical exposures and cardiac disease progression.

In contrast to structural cardiac effects, electrophysiological AOs, such as cardiac arrhythmia or disruption of calcium homeostasis, were comparatively underrepresented in the final network, despite being commonly investigated. This underrepresentation appears to be due primarily to their occurrence at high, often cytotoxic or animal-harming doses, which were excluded during quality filtering. This observation is notable because electrophysiological liabilities are a primary concern in drug-safety evaluation, where hERG blockade, QT-interval prolongation and torsade-de-pointes risk dominate cardiotoxicity screens [18]. The weak signal in our environmentally focused dataset therefore suggests different hazard priorities: While electrophysiological disturbances may be prominent in drug-induced cardiotoxicity, they may be less relevant for environmental chemical exposures at realistic or regulatory-relevant concentrations. This finding also underscores the importance of applying cytotoxicity thresholds (*in vitro*) and well-being criteria (*in vivo*) when interpreting electrophysiological outcomes and highlights the need for endpoint-specific test strategies that reflect the distinct risk profiles of pharmaceuticals versus environmental pollutants.

### Prototypical stressors and modulating factors

To explore whether recurrent mechanistic patterns point to prototypical environmental stressors, we next examined chemical-KE connectivity. Chemicals with extensive study coverage, such as TCDD, cadmium, arsenic, PM₂.₅ and bisphenol A, showed links to multiple high-frequency KEs, notably oxidative stress, inflammation and mitochondrial dysfunction. These associations are consistent with the broader literature. Cadmium and arsenic exposures have been tied to oxidative stress-mediated cardiovascular morbidity in epidemiological and experimental work [19, 20], while PM₂.₅ has been shown to drive cardiac remodelling through inflammatory and redox mechanisms [21, 22]. Notably, the same chemicals identified as key contributors to myocardial abnormalities in our previous epidemiological review, PM₂.₅, NO₂, lead, cadmium, and arsenic, also emerged as mechanistically well-supported stressors in our *in vitro* and *in vivo* evidence mapping. This convergence of epidemiological and mechanistic evidence not only strengthens the case for regulatory attention to these pollutants, but also underscores the validity of our integrative approach for identifying priority stressors and relevant mechanistic pathways in environmental cardiotoxicity.

In contrast, many pesticides, industrial chemicals and engineered nanoparticles were connected to only a handful of KEs and were represented by very few studies, precluding confident assessment of their cardiotoxic potential. This data sparsity may reflect research gaps and publication bias rather than lack of hazard, underscoring the need for targeted mechanistic studies on under-characterised but commonly occurring chemicals. These findings further emphasize the utility of a chemical-agnostic, evidence-based mapping approach: it can highlight well-supported chemical-mechanism relationships while also pointing to substances in need of further investigation to clarify their cardiotoxicity profiles.

Modulating factors such as high-fat diet and pre-existing cardiovascular conditions further indicate that susceptibility and KE intensity are context-dependent, but uneven reporting limits quantitative integration of modifiers into the network at present.

### A consensus, time-resolved AOP network capturing the dominant cardiotoxicity mechanism

Building on the complete set of high-confidence evidence, frequency-filtered KEs, essentiality-supported KERs, and KER-level exposure-duration tags, we distilled an integrated, time-resolved AOP network that represents the central cardiotoxicity mechanisms elicited by environmental chemicals (Figure 7). Based on the most frequently observed and well-supported KEs and KERs, a refined AOP network was constructed to represent the central cardiotoxicity mechanisms triggered by environmental chemicals (Figure 7). This network highlights oxidative stress, mitochondrial dysfunction, and inflammation as core drivers of cardiotoxic outcomes, and traces their progression from acute molecular disturbances through chronic signalling events (pro-hypertrophic and pro-fibrotic events) to chronic tissue remodelling and loss of ventricular function. By annotating every KER with its shortest effective exposure duration, the network provides both mechanistic plausibility and a chronological sequence that can inform *in vitro* assay design and integrated testing strategies. The network illustrates the pathways most consistently reported in the literature and confirmed through our systematic mapping.

**Figure 7:**
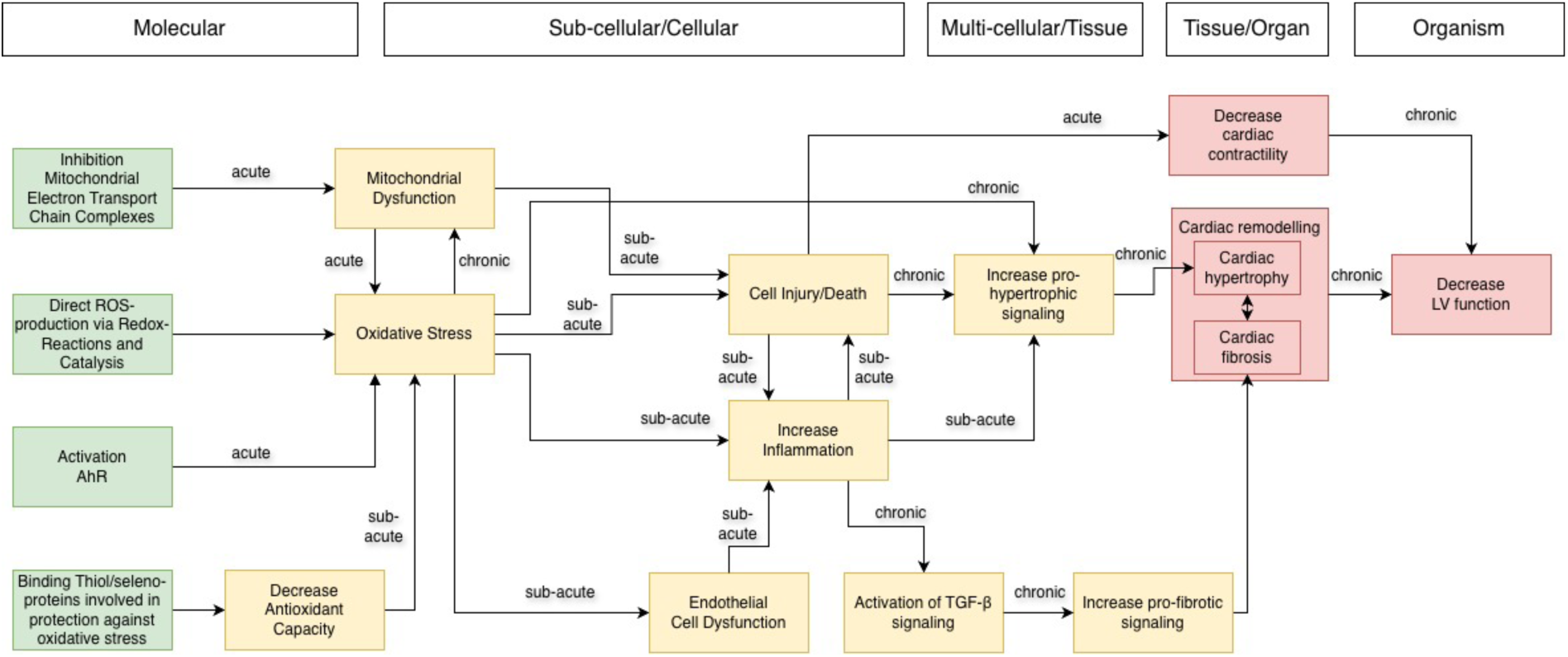
Final time-resolved AOP network for the most frequently observed cardiotoxicity mechanisms induced by environmental chemicals.

While we initially hypothesized that exposure durations might follow a general trend from acute MIEs to chronic AOs, this pattern did not hold when examining KEs in isolation. In fact, several KEs, regardless of their biological level, were observed across a wide range of exposure durations, making temporal interpretation at the KE level inconsistent and potentially misleading, as it ignores the context-specific, relational nature of biological timing. However, when exposure duration was analysed per KER, distinct and interpretable patterns emerged. Each KER, even when involving the same KE, was associated with a characteristic duration that reflected the temporal spacing of the upstream and downstream events. This relationship-level resolution revealed a more accurate picture of cardiotoxic progression, highlighting, for instance, that certain AOs, such as decreased myocardial contractility, can occur after acute exposures, while some upstream or intermediate events (e.g., chronic inflammation or pro-fibrotic signaling) may require subacute or chronic exposure to manifest.

Thus, we extend the AOP concept by embedding an explicit, literature-derived time dimension. Conventional qualitative AOPs list causally linked events but rarely indicate when one event leads to the next, unless quantitative KERs have been modelled. The AOP-Wiki does not yet formally incorporate temporal aspects, such as the duration it takes for one KE to lead to the next or how the timing and duration of exposure (e.g., chronic versus acute) modulate downstream events [23]. Although the scientific community widely acknowledges the importance of temporality for causal interpretation and regulatory application, time is most often discussed in supporting literature or quantitative modelling efforts, rather than embedded in standardized AOP descriptions [23]. Based on our findings, we recommend capturing temporal information, such as whether an event transition is acute or chronic, specifically at the KER level, not at the KE level. This approach directly supports the concept of time concordance, preserves event directionality for regulatory sequence-of-events reasoning, and provides a practical, scalable way to add chronology to the AOP-Wiki without demanding full quantitative AOP modelling.

At the same time, the temporal categorisation remains a pragmatic approximation based on study design windows and does not account for toxicokinetic differences across species or chemicals. As time-course datasets expand and kinetic information becomes more routinely reported, the temporal layer of cardiotoxicity AOP networks can be refined toward more quantitative time concordance.

Methodologically, this work follows a systematic evidence mapping (SEM) approach with transparent, reproducible procedures. A predefined search strategy, explicit inclusion criteria, structured extraction templates, and OHAT-based risk-of-bias filtering ensured comprehensive coverage while maintaining minimum quality standards [24, 25]. This process yielded an evidence-weighted network reflecting the cardiotoxicity mechanisms most consistently reported across the environmental-chemical literature.

## Conclusion

In this study, we developed a bottom-up, literature-derived AOP network for cardiotoxicity induced by environmental chemicals. By translating a large systematic evidence map into harmonised KEs and KERs, applying quality filtering and evidence thresholds, and prioritising mechanistically substantiated relationships via essentiality, we generated a *de novo*, high-confidence mechanistic network that captures the cardiotoxicity pathways most consistently supported across heterogeneous *in vitro* and *in vivo* studies.

A central finding is the convergence of diverse environmental stressors on common stress - cardiac injury events dominated by oxidative stress and mitochondrial dysfunction, which connects to cardiomyocyte injury and progresses toward remodelling outcomes (cardiac hypertrophy and fibrosis) and loss of left ventricular function. Importantly, we demonstrate that temporal information is most informative when represented at the level of KERs rather than KEs: KER-level exposure-duration annotation enables time concordance assessment, improves interpretability of sequence-of-events reasoning, and provides practical guidance for NAM study design by distinguishing rapidly induced functional perturbations from processes requiring sustained exposure.

Overall, this work provides a transparent, evidence-weighted and time-resolved cardiotoxicity AOP network that can support the development of cardiotoxicity-focused NAM batteries, guide prioritisation of endpoints and exposure durations, and strengthen mechanistically informed interpretation of environmental chemical hazards. Future efforts should expand coverage of MIEs, improve consistency in reporting time-course data, and further refine temporal annotation as additional longitudinal mechanistic datasets become available.

## Author contributions

Alexandra Schaffert, Sivakumar Murugadoss, and Tom Roos contributed equally and share first authorship. Birgit Mertens and Martin Paparella contributed equally and share last authorship. Alexandra Schaffert, Sivakumar Murugadoss, Tom Roos, Birgit Mertens, and Martin Paparella conceived the study and designed the methodology. Birgit Mertens, and Martin Paparella provided project resources. Alexandra Schaffert, Sivakumar Murugadoss, and Tom Roos conducted data extraction. Alexandra Schaffert led data curation and analysis. Nunzia Linzalone, Gabriele Donzelli, and Ronette Gehring contributed to methodology design, and evidence interpretation. Alexandra Schaffert, Sivakumar Murugadoss, and Tom Roos drafted the manuscript. All authors reviewed, edited, and approved the final manuscript.

