## Supplementary 1 for "A Data-Driven Approach for the Development of a Time-informed Adverse Outcome Pathway-network for Cardiotoxicity of Environmental Chemicals"

Table S1: Species- and system-specific reference exposure durations and classification thresholds used for exposure time categorization.

| **Experiment Type** | **Species category** | **Acute ≤** | **Subacute >…≤** | **Subchronic >…≤** | **Chronic >** | **Acute requires single dose?** | **Short continuous allowed as Acute?** | **Justification** |
| --- | --- | --- | --- | --- | --- | --- | --- | --- |
| in_vivo | Mice | 1 d | 28 d | 90 d | 90 d | Yes | No | OECD acute oral TGs are single-dose with 14-d observation; 28- and 90-day repeated-dose define subacute/ subchronic (TGs 420, 423, 407, 408) |
| in_vivo | Rats | 1 d | 28 d | 90 d | 90 d | Yes | No | Same rationale: acute = single dose; TG 407 (28 d) and TG 408 (90 d) for longer windows |
| in_vivo | Zebrafish (incl. embryos) | 4 d | 21 d | 60 d | 60 d | No | Yes | Aquatic “acute” tests are 24–96 h (fish acute TG 203; FET TG 236); ELS TG 210 extends to early juvenile → places 21–60 d bounds |
| in_vivo | Other: Drosophila melanogaster | 4 d | 21 d | 60 d | 60 d | No | Yes | No OECD TG; ~10 d generation time and ~60–70 d lab lifespan justify ≤4 d “acute” and >21–60 d subchronic bounds [1] |
| in_vivo | Swine (incl. pigs) | 1 d | 28 d | 91 d | 91 d | Yes | No | Non-rodent general tox practice per ICH M3(R2): ~13 wk subchronic; ≥6–9 mo chronic. We set the boundary at 13 wk |
| in_vivo | Chicken (incl. broilers & eggs) | 1 d | 70 d | 140 d | 140 d | Yes | No | Acute oral TG 223 = single dose with 14-d observation; avian reproduction TG 206 collects eggs for ~10 wk = 70 d, anchoring higher categories |
| in_vivo | Rabbits | 1 d | 28 d | 91 d | 91 d | Yes | No | Treat as non-rodent per ICH M3(R2) (subchronic ≈13 wk; chronic ≥6–9 mo) |
| in_vivo | Other: Tadpoles | 4 d | 21 d | 60 d | 60 d | No | Yes | Amphibian Metamorphosis Assay TG 231 is a 21-day exposure; ≤4 d follows aquatic “acute” convention; >21–60 d subchronic; >60 d chronic |
| in_vivo | Quails (incl. eggs) | 1 d | 70 d | 140 d | 140 d | Yes | No | Same avian anchors as chicken: TG 223 (single dose + 14 d obs) and TG 206 (~10 wk egg-collection) |
| in_vivo | Other: Marmosets | 1 d | 28 d | 91 d | 91 d | Yes | No | Non-rodent primate durations per ICH M3(R2); many chronic studies are 26–52 wk |
| in_vitro | all species | 3 d | 14 d | 28 d | 28 d | No | Yes | Validated assays use 24–72 h exposures (e.g., KeratinoSens™ 48 h TG 442D; h-CLAT ~24 h TG 442E). GIVIMP frames exposure as method-defined |

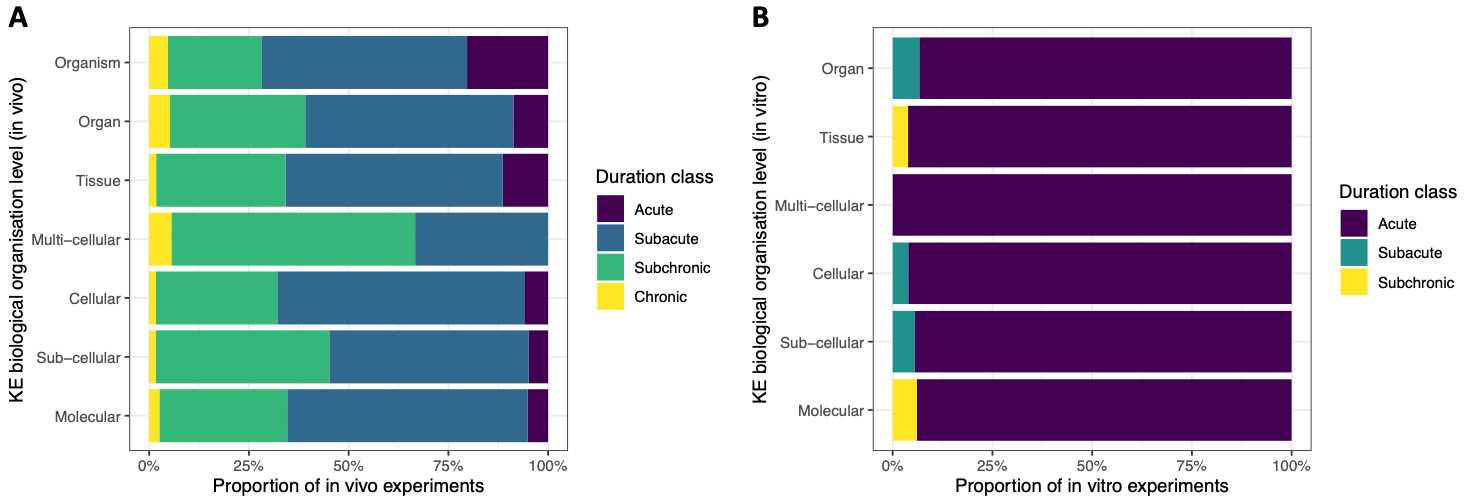

Figure S1: Duration classes for in vivo (A) and in vitro (B) KEs on different biological organisation levels.

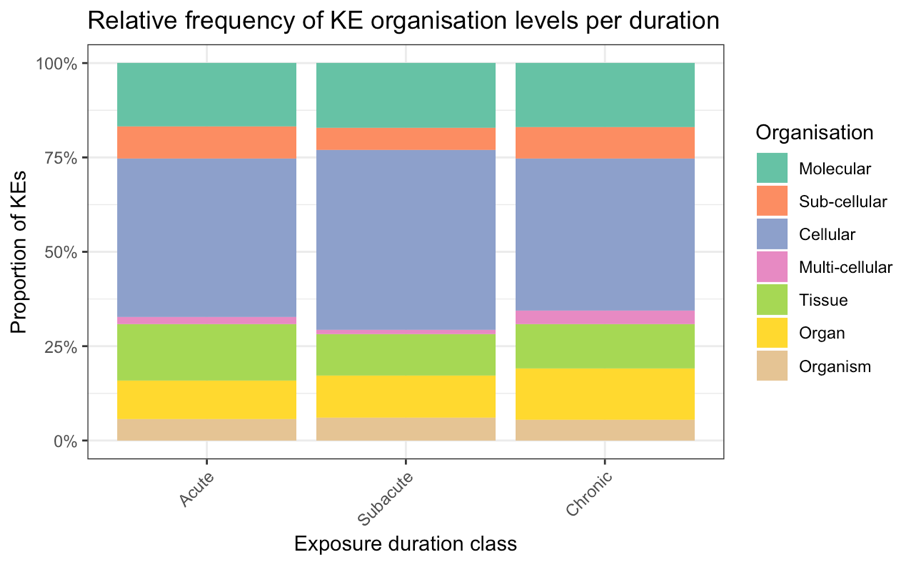

Figure S2: Reltie frequency of KE organisation levels per duration.

1.	Ogienko, A.A., et al., *Drosophila as a Model Organism to Study Basic Mechanisms of Longevity.* Int J Mol Sci, 2022. **23**(19).
